# Learning the bistable cortical dynamics of the sleep-onset period

**DOI:** 10.1101/2025.07.17.665340

**Authors:** Zhenxing Hu, Manaoj Aravind, Xu Lei, J. Nathan Kutz, JJ Aucouturier

## Abstract

Humans just don’t fall asleep like a log – or step-function. Rather, the sleep-onset period (SOP) exhibits dynamic and non-monotonous changes of electroencephalogram (EEG) with high, and so far poorly understood, intra- and inter-individual variability. Computational models of the sleep regulation network have suggested that the transition to sleep can be viewed as a noisy bifurcation at a saddle point which is determined by an underlying control signal or “sleep drive”. However, such models do not describe how internal control signals in the SOP can produce rapid switches between stable wake and sleep states, nor how these state-space changes are translated in the macroscopic EEG. Here, we propose a minimally-parameterized stochastic dynamical model, in which one slowly-varying control parameter drives the wake-to-sleep transition while exhibiting noise-driven bistability. We provide a procedure for estimating the parameters of the model given single observations of experimental sleep EEG data, and show that it can reproduce a wide variety of SOP phenomenology. Using the model to analyze a pre-existing sleep EEG dataset, we find that the estimated model parameters correlate with subjective sleepiness reports. These results suggest that the bistable characteristics of the SOP can serve as biomarkers for tracking intra- and inter-individual variability of sleep-onset disorders.

**Author summary:** Recent neuroscience research has showed a growing interest in understanding the complexity of how we fall asleep. Electroencephalographic (EEG) recordings of the sleep onset period show all the hallmarks of a noise-driven bistable system, but there currently exists no computational model that can be fitted on experimental data to understand this behavior. In this paper, we propose a minimally-parameterized model, which dynamics corresponds to the motion of an noisy overdamped particle in a slowly tilting bistable landscape, as well as a way to fit it to an individual’s sleep-onset EEG. We show that the fitted parameters of individual participants correlate with their subjective reports of sleepiness, suggesting that the model can capture important aspects of inter-individual variability, as well as provide potential biomarkers for sleep-onset disorders.

## Introduction

The ability of organisms to keep track of the time of day and maintain cycles of stable wake and sleep states has fascinated physiologists for a large part of the 20th century [1], and has become an iconic target of research for mathematical and dynamical-system modeling [2]. Following seminal work by Borbély, Daan and Beersma [3], mathematical models for sleep-wake regulation have traditionally included the interaction between two coupled processes: a relaxation oscillator (the homeostatic drive) by which ‘sleep pressure’ monotonically increases during wake and relaxes during sleep; and a circadian oscillator which modulates homeostatic thresholds approximately sinusoidally [4]. While phenomenological ^1^ in nature, these models were found in good accordance with predictions made by more complex biophysical models of the ascending arousal system, such as Phillips’ and Robinson’s [6], and have been used to explain such diverse phenomena as sleep restriction experiments, the effect of caffeine, or changes in sleep patterns during development (for a review, see [7]).

Among other fascinating sleep-related dynamical phenomena, the transitional phase between wake and sleep (‘sleep-onset period’, or SOP) has attracted recent attention in the neuroscience community [8]. While long regarded as a monotonic process akin to “ flicking a switch” or, alternatively, a long sequence of successive substates [9], the SOP is now widely regarded as a continuous, dynamic process that fluctuates progressively, but non-monotonically, between wake and sleep, and with high heterogeneity both within and between individuals [10]. At the surface EEG level, the wake-to-sleep transition is primarily marked by the progressive disappearance of the EEG alpha rhythm, and exhibits all the hallmarks of bistable behaviour (Fig. 1-top). However, while it is increasingly suspected that the internal dynamics of the SOP has both cognitive and clinical significance in subsequent sleep and wake periods [11, 12], there exists little mechanistic understanding of what produces such patterns and their heterogeneity. Even at a basic phenomenological level, the SOP and its associated first stage of sleep (N1) remains the period with the lowest inter-scorer agreement and classification accuracy in both humans [13] and machines [14].

**Fig 1.**
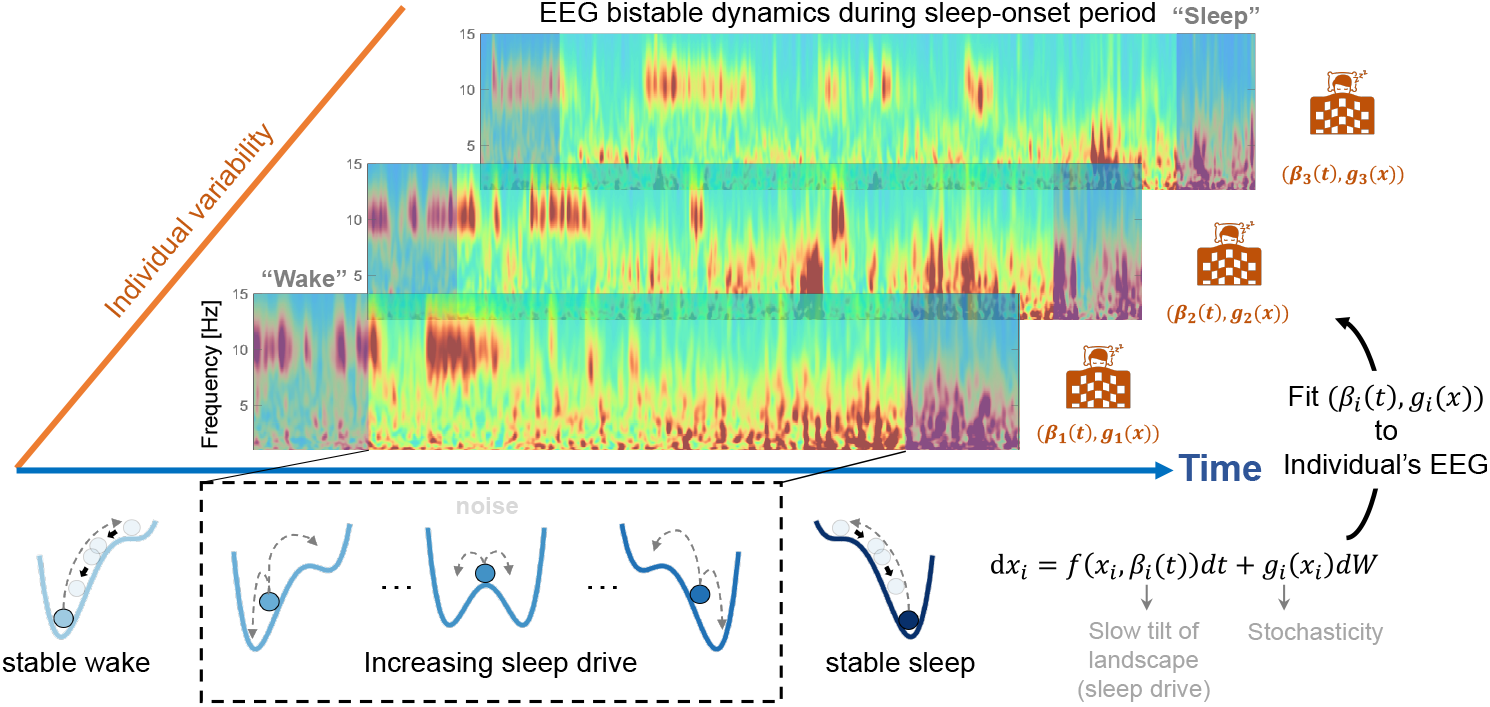
Study overview. The sleep-onset period (SOP) has a strongly bistable phenomenology, marked by a non-monotonous decrease of the EEG frequency and high inter-individual variability, seen here in three illustrative wavelet spectrograms (**top**). We model the bistable cortical dynamics of the SOP by introducing a minimally-parameterized stochastic dynamical system, which dynamics corresponds to the motion of an noisy overdamped particle in a slowly tilting bistable landscape (**bottom**). We provide a procedure to estimate model parameters given individual observations of experimental sleep EEG data (**right**), which allows us to test whether model parameters correlate with clinical feature.

In recent work [15], the Philipps-Robinson (PR) model of the all-day sleep regulation network [6] was used to approximate the dynamics near the transition from wake to sleep. At the tipping point corresponding to the critical sleep drive value necessary for the transition to occur, the dynamics can be reduced to the equation of motion of a particle in a frictional and approximately cubic potential. In this framework, the SOP is, therefore, modeled as a saddle-point bifurcation between stable wake and stable sleep, which the authors propose to be driven by gradual increase of the sleep drive, which comprises the approximately 24-h periodic circadian drive from the suprachiasmatic nucleus (SCN) and the homeostatic drive due to sleep homeostat (HOM). While their reduced model could generate analytical scalings of fluctuation variance and spectral width in state-space, that is consistent with numerical simulation by the full PR model, it remains consistent with a traditional, ‘monotonic’ view of the SOP, and fails to describe how internal control signals (or noise) during the SOP can produce the back-and-forth switches between wake and sleep that are seen in real experimental data.

In addition, this and other related stochastic models of wake-sleep transition behaviour [16, 17] only provide simulations of the phenomenon of interest (‘how possibly’ explanations in the sense of [18]), but do not describe how these state-space phenomena are translated in the macroscopic EEG. First, they do not provide a data-driven embedding of the (high-dimensional, measurement-noise prone) EEG data in which these simplified dynamics are effected. Second, they do not provide a fitting procedure by which model parameters can be identified given experimental sleep data - a non-trivial task when the underlying models are stochastic and non-stationary, and data may come in only one single observation per participant (i.e. one given SOP). In the absence of such data-driven procedure, SOP models fall short of explaining the occurrence of a particular EEG pattern and, perhaps more importantly, their association with concurring disorders such as insomnia or anxiety in specific individuals.

To do so, we propose a minimally parameterized stochastic dynamical model in which the system’s state is governed by a bistable potential landscape (Fig 1, see *Materials and Methods* for details). In previous work, a similar form was used to model how external drivers control switches between the two neural states that correspond to forward and backward motion in C. Elegans [19]. Here, the system’s state is assumed to move on a continuum between two potential basins correspond to the ‘wake’ and ‘sleep’ states and, instead of using an external control signal to actuate the state into sleep, we let a putative ‘sleep drive’ tilt the landscape towards the sleep basin (Fig 1, left to right), with the result of making noise-driven transitions to sleep not inevitable, but increasingly more likely, over the SOP.

In addition, we construct a generalized, low-dimensional representation of the SOP EEG by performing a low-rank SVD decomposition of the EEG spectrogram and use the linear interpolation between the two wake and sleep SVD modes to represent the dynamics linked to the transition. We provide and validate a procedure for fitting our dynamical system model to the trajectory of that interpolation, and show that its dynamics can reproduce a wide-variety of SOP phenomenology. Finally, using the model to analyze a preexisting sleep (nap) EEG dataset, we test whether the estimated model parameters correlate with subjective sleepiness reports collected around the nap.

## Results

### The bistable characteristics of the SOP are preserved in a low-rank embedding of the EEG spectrogram

Sleep-onset periods (SOP) in our EEG dataset of healthy adults have a typical, strongly bistable phenomenology, marked by a non-monotonous decrease of the EEG alpha (8-12Hz) frequency [8]. In a typical participant (Fig. 2a-top), the alpha component exhibits a prolonged, relatively stable amplitude early in the analysis window (here, from about 0s to 100s). This alpha activity gradually becomes more transient and intermittently suppressed, indicative of a “bistable” pattern in which the signal switches between high and low amplitude on a short timescale. In the transitional period (inside the dashed line), there are rapid switches between alpha component and the low-frequency component. Beyond 300s, the alpha band is largely diminished and replaced by an increasingly dominant low-frequency (0.5–4 Hz) component, reflecting the transition from wake into sleep-like regimes.

**Fig 2.**
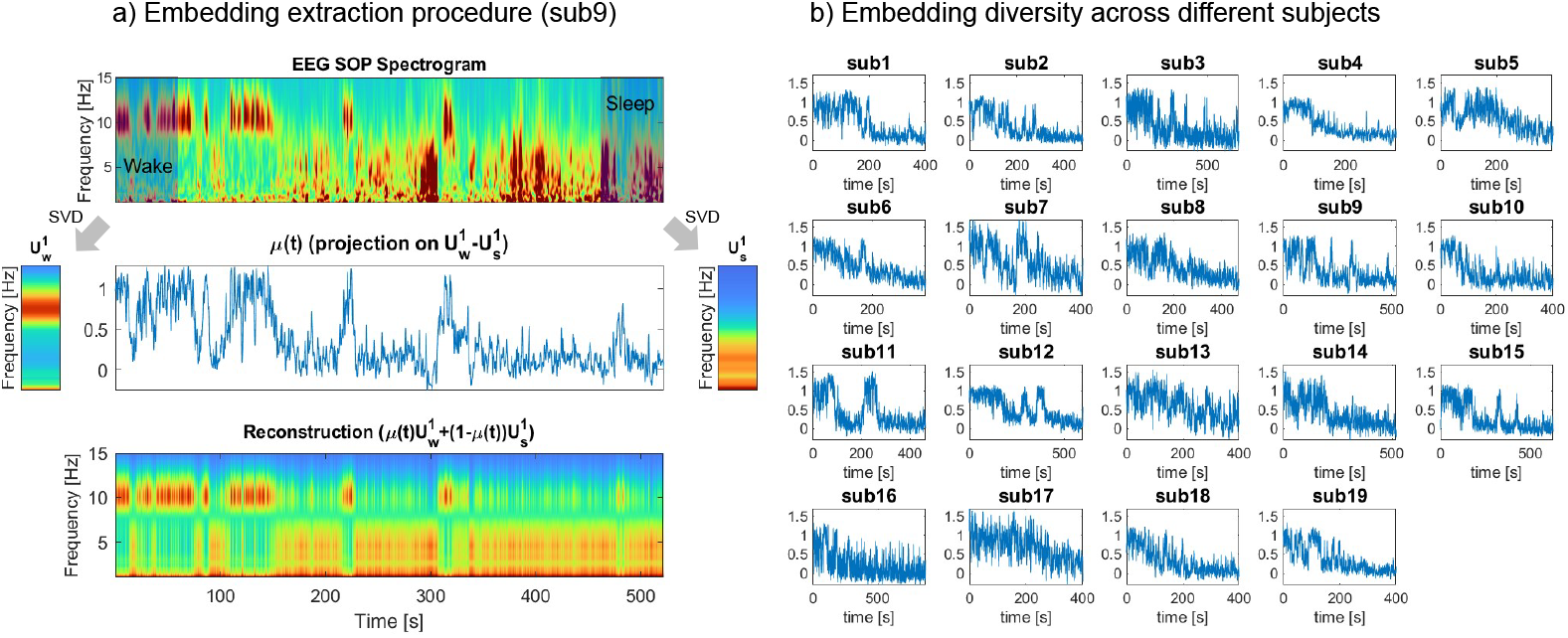
Bistable dynamics are manifested in a low-rank embedding of the SOP EEG spectrogram. (a) single-subject example. **Top:** Spectrogram representation of a single-channel (Oz, occipital, median) EEG recording of the SOP, from wake (left) to sleep (right), for one illustrative participant. Grayed windows identify the initial stable wake and final stable sleep phases, and dashed lines represent the start and the end of transitional period. **Middle:** Dominant SVD modes for the wake (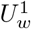 , **left**) and sleep states (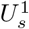, **right**) extracted from the initial and final windows. A low-dimensional representation *µ*(*t*) is obtained by projecting the normalized spectrogram to the principal direction 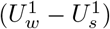, and captures the major transitions and shifts from wakefulness to sleep. **Bottom:** Reconstructed spectrogram obtained from the *µ*(*t*) embedding by linear interpolation between the wake and sleep mode. **(b)** *µ*(*t*) embeddings extracted from N=19 healthy participants in our test dataset (see *Materials and Methods*).

We construct a generalized, low-dimensional representation of the SOP spectrogram by computing the singular value decomposition (SVD) of the spectrogram separately in the initial wake and final sleep segment (see *Materials and Methods*), with dominant modes 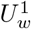 and 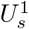 respectively (Fig.2a-middle). We then generate an embedding *µ*(*t*) of the SOP spectrogram by projecting the spectrogram on 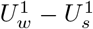. Coefficient *µ*(*t*) maps how strongly the EEG at each moment aligns with the wake versus sleep mode.

This coefficient neatly compresses the broad changes seen in the high-dimensional time-frequency plot - including shifts in dominant frequency modulation - into a one-dimensional time series (Fig.2a-middle).

This low-dimensional representation provides a state-space for our model. Specifically, we fit below our general dynamical systems model to the trajectories of the *µ*(*t*) embedding. Conversely, the embedding also allows us to project state-space dynamics back into observation (EEG) space, using the linear interpolation 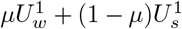 (Fig. 2a-bottom). Although some finer details are inevitably lost in this low-dimensional representation, the major transitions and overall shifts from wakefulness to sleep are captured by this low-rank approximation. It also captures important inter-individual variations (Fig. 2-b), which motivates the need to fit model parameters at the individual level and look for potential associations of these parameters with individual sleep characteristics.

### Changes to two model parameters reproduce a wide variety of SOP phenomenology

To model the dynamics of *µ*(*t*), we propose a minimally parameterized first-order model of the form,

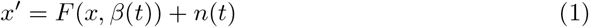

where *F* (*x, β*(*t*)) represents the intrinsic nonlinear dynamics containing two stable states separated by a saddle; *β*(*t*) is a control parameter dictating the shape of landscape that evolves over long time scales, and *n*(*t*) is additive noise adding stochastic forcing to the intrinsic dynamics. The nonlinear dynamics *F* (*x, β*(*t*)) is modeled by the cubic function

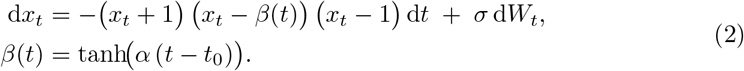

which has two stable fixed points at *x* = *±*1, and a single unstable fixed point at a time-varying location *β*(*t*) which is parameterized by a slowly-drifting hyperbolic tangent function with slope *α* and offset timing *t*_0_ (Fig 3). In broader context, the phenomenon of nonlinear interactions between neighboring stable (fixed) points is also a well-studied representative model in optical and atomic physics, with nonlinearity producing nontrivial dynamics in the dynamics [20, 21]. The cubic model considered is also a canonical model for studying the Langevin dynamics between two stable states [22]. Thus (2) is a canonical model for many systems where the nonlinear dynamics between two stable states drives the observed transition phenomenon.

**Fig 3.**
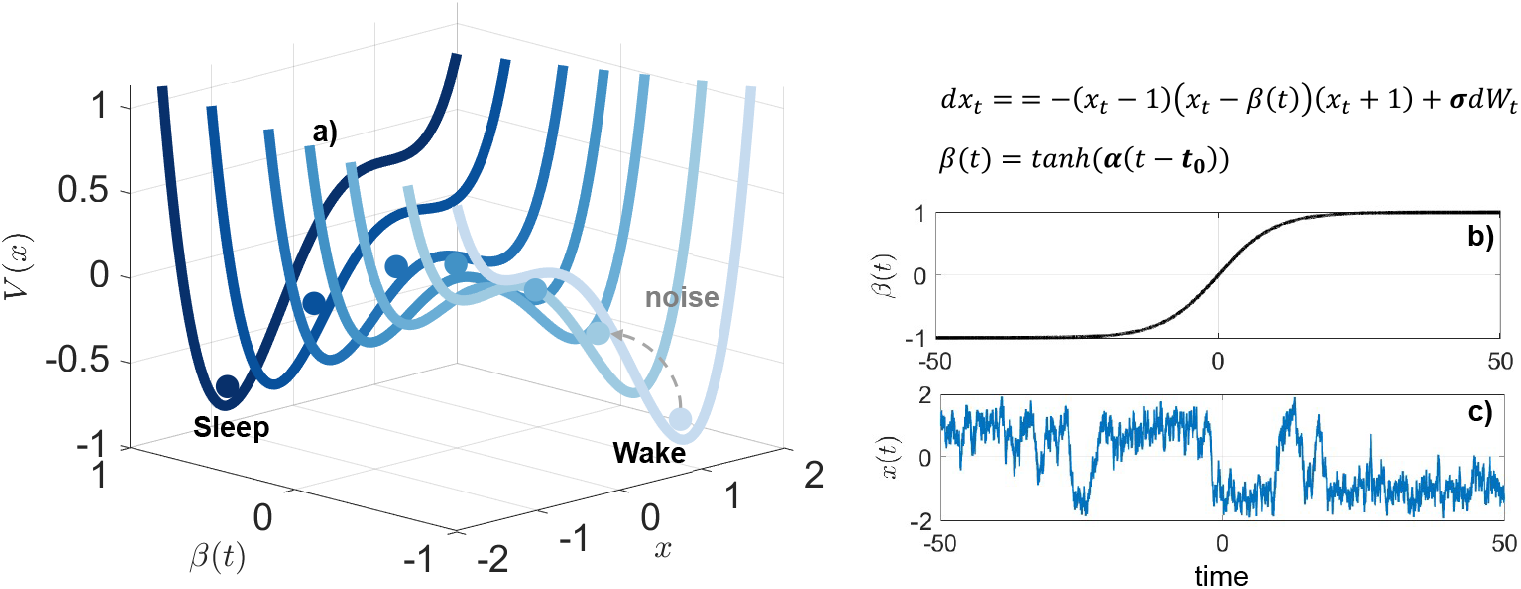
Schematics of time-varying stochastic bistable model. a) The potential functions, *V* (*x*), transitions from a “wake” basin (right) to a “sleep” basin (left) as *β* shifts from -1 to +1. b) *β*(*t*) = *tanh*(*α*(*t* − *t*_0_)) governs the gradual tilt of the landscape over time. c) A representative trajectory *x*(*t*) simulated from the stochastic differential equation Eq. 2).

By controlling how quickly the landscape changes (*α*) and how strongly the dynamics is subjected to stochasticity (*σ*), the model can capture a range of transition behaviors, from monotonous gradual shifts to abrupt, noise-driven back-and-forth switches (Fig 4). In general terms, larger *α* values tend to induce earlier transitions to sleep (compare Fig 4-top and middle rows), and larger *σ* values lead to increased flickering between the wake and sleep states before settling into one (compare Fig 4-middle and right columns). However, the influence of the two parameters is not entirely independent, nor linear.

**Fig 4.**
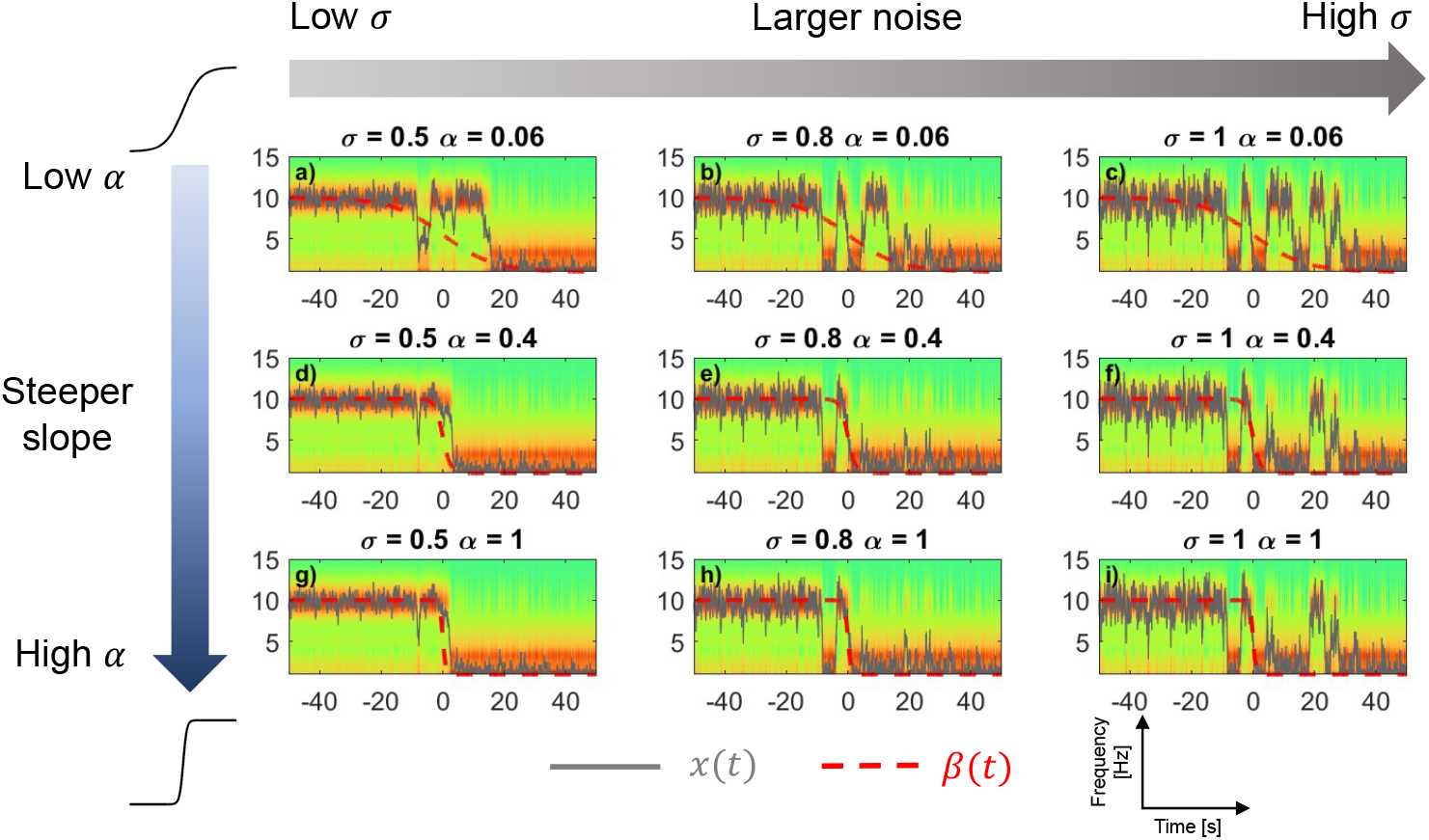
Interactions between the rate of landscape change (*α*) and noise (*σ*) in wake-to-sleep transitions. Each panel shows a simulated spectrogram over time, with *α* increasing from top to bottom (0.06, 0.4, 1) and *σ* increasing from left to right (0.5,0.8,1). In general terms, larger *α* values tend to induce earlier transitions to sleep (compare top and middle rows), and larger *σ* values lead to increased flickering (compare middle and right columns). All *x*(*t*) (grey) are normally stochastic, but simulated here with the same random seed for the purpose of comparison. *β*(*t*) (red, dashed) is normalized in amplitude, also for visualization. EEG embedding similar as Fig. 2.

First, as *α* grows, the model exhibits a saturation effect in which, after a certain point, transition to sleep does not occur earlier for larger values of *α* (Fig 4-left column, middle and bottom rows). This is due (i) to the nonlinear relationship between *α* and the timescale of the underlying *tanh* function, and possibly (ii) to sampling-rate limits of the dynamical system itself, which cannot track very rapid landscape shifts beyond a certain threshold. Second, at fixed levels of *α* (e.g. Fig 4-top row), increasing *σ* level can induce faster transitions. Finally, at large *α* values, the pronounced “tilt” in the landscape can keep the system in the sleep state after transitioning, even when higher noise levels are present. The wide variety of this phenomenology in state-space can be explained by the interplay between the landscape and stochasticity, which potentially lead to the empirical inter-individual variations seen in Fig 2-b.

### Model parameters can be recovered from a single experimental trajectory of the system

Infering likely values for parameters *α, t*_0_ and *σ* given a single realization, i.e. one participant’s SOP EEG spectrogram is made difficult because the model is (1) stochastic (so a single value of *σ* can lead to an infinite number of realizations) and (2) non-stationary (so a given intermediate value *β*_*i*_ = *β*(*t*_*i*_) can only be fitted given one time sample of the observed trajectory). In Section *Materials and Methods*, we provide a Markov Chain Monte Carlo (MCMC) formulation of the parameter-fitting problem. Here, we use simulated data to evaluate how the procedure recovers the parameters under both stationary and non-stationary assumptions, and then illustrate parameter estimation with one representative example of experimental SOP data.

#### Stationary landscape

when the double-well potential is stationary (i.e., *β*(*t*) = *β*), the system’s trajectory only reflects noise-driven fluctuations within a fixed landscape. Under these assumptions, we simulated 30 random realizations (*x*(*t*) trajectories) for every pair of values *β* ∈ [−0.8, 0.8] and *σ* ∈ {0.2, 0.5, 1.0} and evaluated recovered parameters for each trajectory with MCM. The procedure was able to recover both *β* and *σ*. For *β*, posterior-mean estimates over the 30 trajectories showed good agreement with the true *β* values (Fig 5-left), but the standard deviation over individual trajectories, although relatively moderate, was non-negligible, suggesting that some degree of degeneracy (i.e. different *β* values may yield similar trajectories). For *σ*, MCMC estimates closely tracked true values (Fig 5-right), showing both low bias and low variance. This likely reflects that the underlying likelihood function is more sensitive to changes in *σ*, making noise-related parameters easier to pin down compared to the possibly overlapping trajectories that can occur for varying *β*.

**Fig 5.**
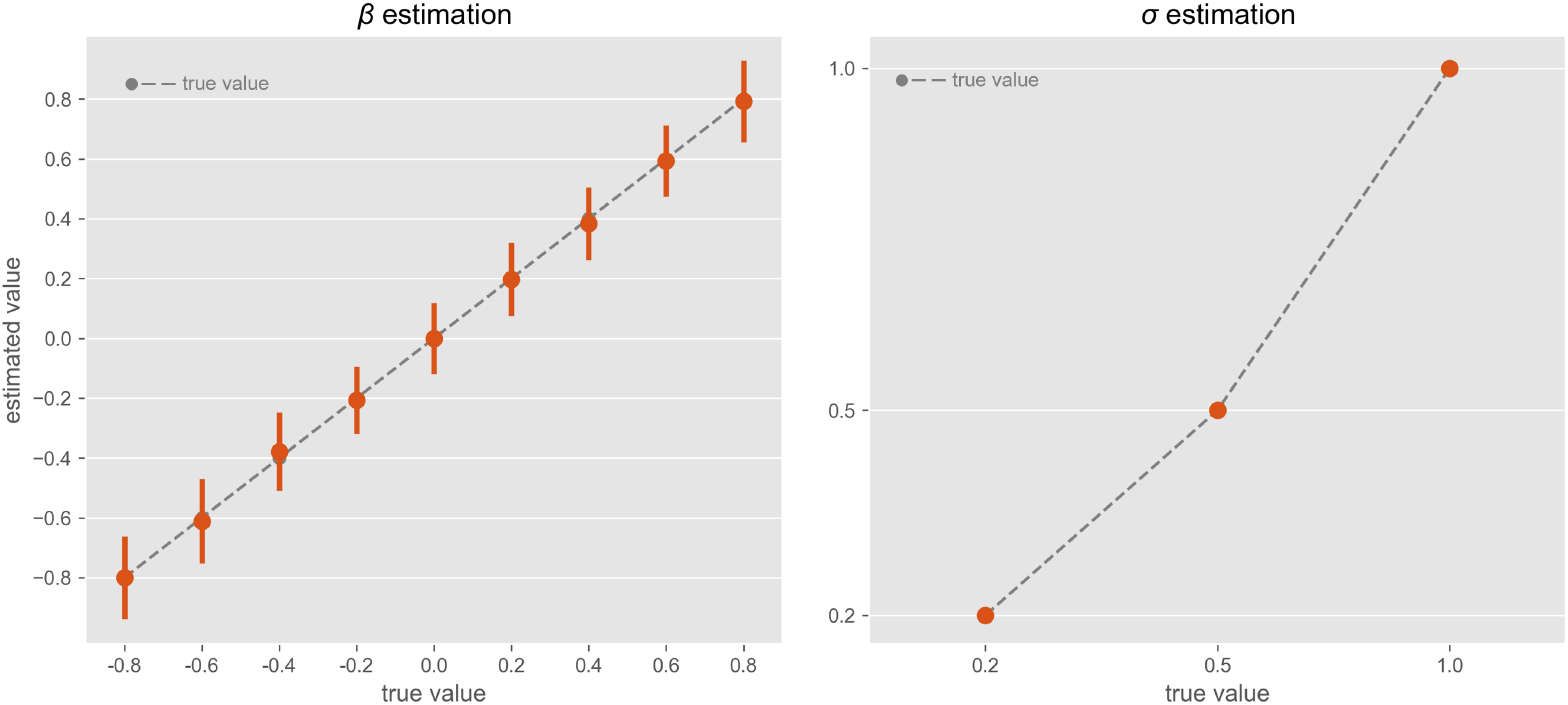
Model parameter estimation for simulated single trajectories under the stationary landscape assumption (*β*(*t*) = *β*). **Left:** 30 trajectories were simulated for every value of *β* ∈ [−0.8, 0.8]. Distributions of MCMC estimates for *β* (y-axis) are plotted against the true values (x-axis). Error bars indicate standard deviations of estimates across multiple trajectories of the same *β* value. **Right:** 30 trajectories were simulated for every *σ* ∈ *{*0.2, 0.5, 1.0}. Distributions of MCMC estimates for *σ* (y-axis) are plotted against the true values (x-axis). Error bars are barely visible, but similarly indicate one standard deviations across multiple trajectories of the same *σ*.

#### Non-stationary landscape

In scenarios with a time-varying landscape, the procedure needs to estimate not only noise-level *σ* but also *t*_0_ (the tipping point at which the potential starts to change) and *α* (the steepness of that change). As before, we simulated 30 random trajectories for every value of *α* ∈ [0.05, 3] (logarithmically-spaced), *t*_0_ 4, 5, 6 and and *σ* ∈ {0.2, 0.5, 1.0} , and evaluated recovered parameters with MCMC estimation.

As in the fixed-landscape scenario, parameter *σ* was accurately recovered for each individual simulation (Fig 6-b) with minimal standard deviation across multiple realizations of the same noise level (Fig 6-b). Also, *t*_0_ estimation (Fig 6-a) was largely accurate, only deviate slightly upward when the true *t*_0_ = 40. This bias is possibly due to the normal prior centered at *t*_0_ = 50, as well as less sensitivity of likelihood to changes of *t*_0_. Despite this bias, the ordering of the estimated *t*_0_ still reflects the true ordering (i.e., the rank relationship is preserved), which is sufficient for distinguishing earlier from later transitions. Similarly for *α*(Fig 6-c), the MCMC procedure overestimated small and underestimated large *α* values, although it preserved ordering (Spearman r = 0.996 across whole range of *α* values). Notably, as *α* increases beyond approximately 1 – 1.5, the estimated values begin to plateau, indicating a saturation effect. This is possibly because, as *α* becomes large, the slope in the underlying *tanh* function saturates (Fig 6-d) due to the inverse relationship between the time-scale of *β*(*t*) and *α*, plausibly making it more difficult for the estimation procedure to distinguish further increases in *α*.

**Fig 6.**
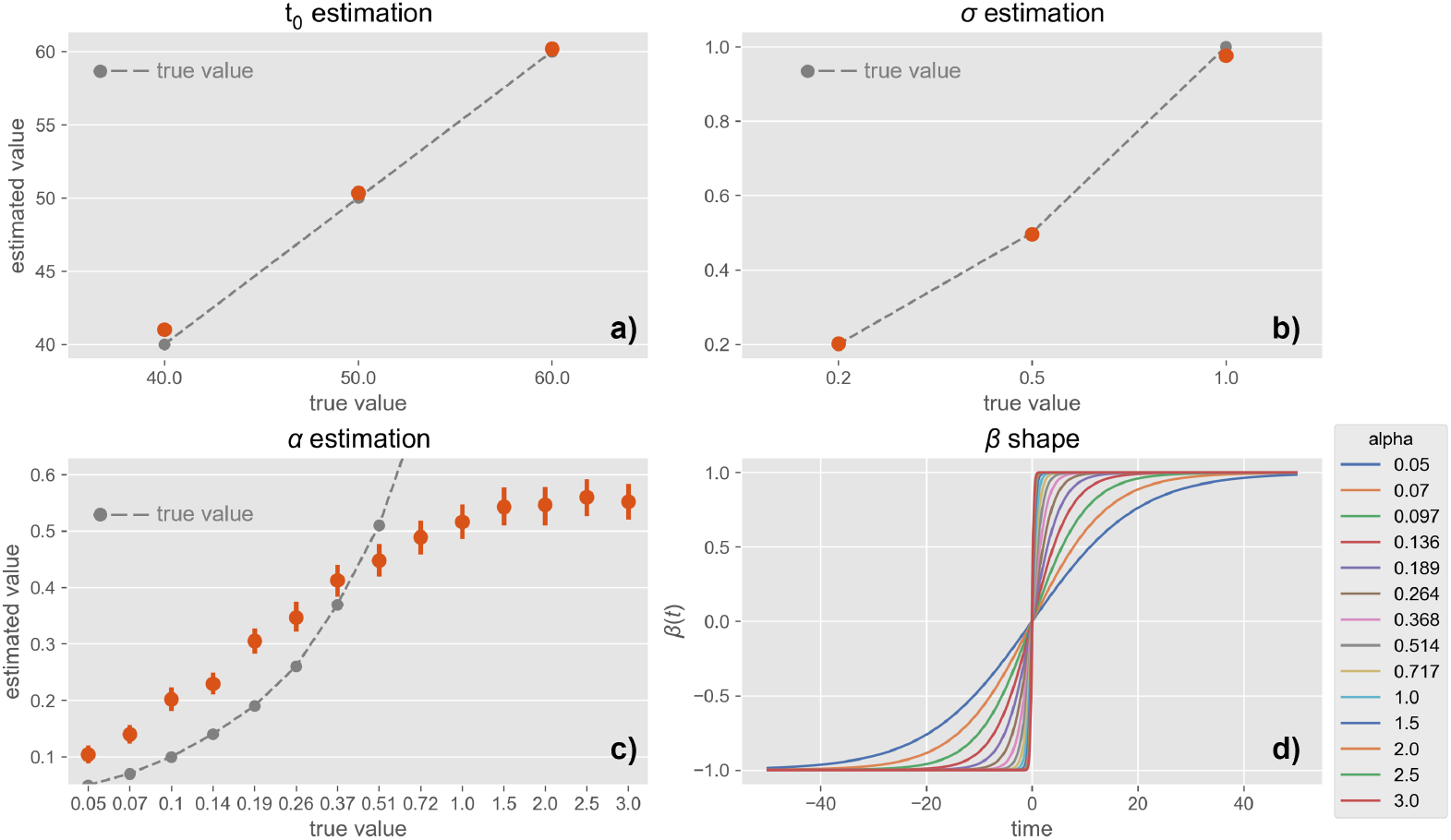
Model parameter estimation for simulated single trajectories under the varying-landscape, non-stationary assumption. **A:** 30 trajectories were simulated for every value of *t*_0_ ∈ {40, 50, 60} and recovered MCMC estimates for *t*_0_ (y-axis) are plotted against true values (x-axis). Error bars (barely visible) indicate standard deviations across multiple realizations of the same *t*_0_, and dashed line indicates perfect estimation. **C**. 30 trajectories were simulated for every value of *σ* ∈ {0.2, 0.5, 1.0}, and MCMC estimates (y-axis) are plotted against true values (x-axis). Error bars (barely visible) indicate standard deviations across multiple realizations of the same *σ*, and dashed line indicates perfect estimation. **C:** 30 trajectories were simulated for every value of *α* ∈ [0.05, 3.0] (logarithmically-scaled), and MCMC estimates (y-axis) are plotted against true values (x-axis). Error bars indicate standard deviations across multiple realizations of the same *σ*, and dashed curve indicates perfect estimation. **D:** Varying *β*(*t*) shape (x-axis:time) as a function of slope *α*, illustrating a saturation effect at larger *α* values.

#### Validation with experimental data

Finally, we illustrate here parameter estimation, and the associated reconstructed EEG spectrogram on one illustrative example of experimental SOP data, recorded from one participant (female, 19; see *Materials and Methods*).

To compare the fitted model with the observed data, we used posterior predictive check by generating 4000 trajectories from the joint posterior distribution over parameter sets, and computed two metrics for goodness of fit: Kullback-Leiber (KL) divergence, which measures how closely the statistical distribution of a single simulated trajectory resembles that of the real EEG embedding, and Root Mean Square Error (RMSE), which reflects point-wise temporal alignment between model output and data. Neither metric alone is perfect; KL divergence ignores temporal correlations in time, whereas RMSE does not capture model stochasticity due to its point-wise nature.

We display the distribution of both metric scores (Fig 7-a). The distribution of KL-Divergence and RMSE over 4000 simulated trajectories of the estimated models shows that most KL values cluster near 0.16, and RMSE around 0.87, indicating that for many model realizations from the posterior point clouds, the distribution of states is moderately close to the empirical data. The sampled trajectory with minimal KL-distance to the true trajectory over the set of 4000 trajectories is obtained for KL = 0.06. Visually, this trajectory has a similar distribution of *x*(*t*) to the original embedding (Fig 7-b), confirming that the minimal-KL solution indeed approximates the overall dwell-time distribution of the embedding *µ*(*t*). The trajectory corresponding to minimal RMSE *x*_*min RMSE*_(*t*) over the sampled set of trajectories aligns relatively well with the original *µ*(*t*) (Fig 7-c), and the real embedding remains with the 10%-90% range of trajectories simulated from the joint posterior distribution. The corresponding reconstructed spectrogram in Fig 7-g can be compared with the original SVD embedding in Fig 7-f, confirming the model’s ability to generate realistic spectrograms.

**Fig 7.**
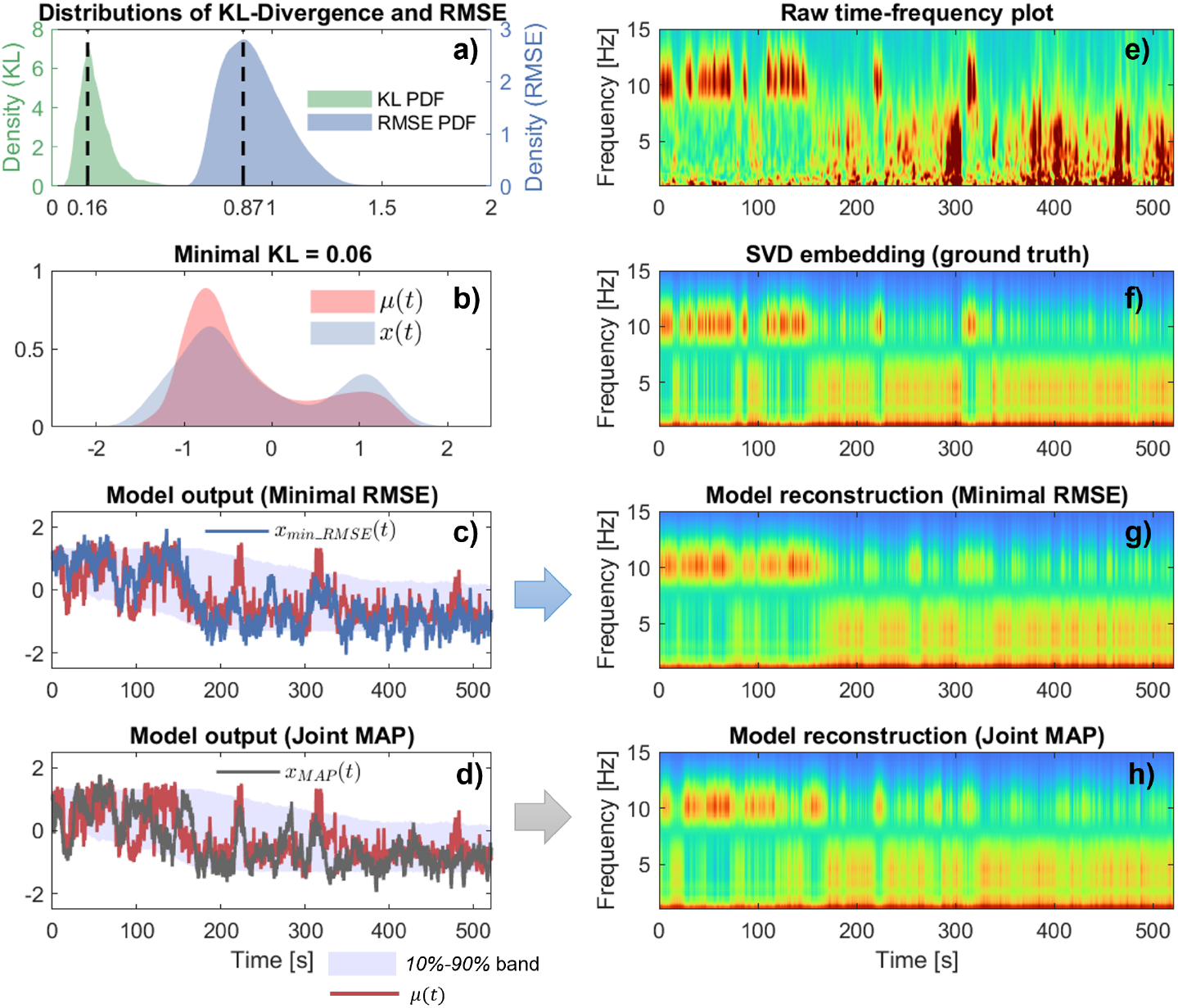
Illustration of fitted parameters and model reconstruction on experimental EEG data. **(a)** Distributions of the Kullback–Leibler (KL) divergence and Root Mean Sqaure Error (RMSE) between the original trajectory and 4000 simulated trajectories from the posterior distributions of estimated model. Solid lines identify the simulated trajectories with the mode KL, and mode RMSE, over the random set. **(b)** Comparison of the probability density function of the minimum-KL trajectory (blue) with that of the original trajectory *µ*(*t*) (red). **(c)** Time-domain comparison of the minimal-RMSE solution (blue) with the original embedding *µ*(*t*) (red). Shaded areas correspond to the 10% - 90% amplitude span of solutions from the 4000 sampled trajectories. **(d)** Time-domain comparison of the Joint MAP solution (dark gray) with the original embedding *µ*(*t*)(red). Shaded areas correspond to the 10% - 90% amplitude span of solutions from the 4000 sampled trajectories. **(e)** Wavelet spectrogram of the original participant EEG. **(f)** Reconstructed spectrogram from the original *µ*(*t*) embedding, obtained using the linear interpolation 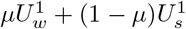 (similar to Fig. 2a)-bottom). **(g)** Reconstructed spectrogram from the minimum-RMSE solution of panel c. **(h)** Reconstructed spectrogram from Joint MAP solution of panel d.

Similarly, the simulated trajectory with joint maximal posterior density (joint MAP) to the experimental data *x*_*min RMSE*_(*t*) is illustrated in Fig 7-d, and the corresponding spectrogram reconstruction in Fig 7-h. The Joint MAP reconstruction replicates global spectral transitions and captures the intermittent switching phenomenon. While the trajectory does not match point-by-point, due to the stochastic nature of the model, it captures the bistable dynamics and eventual transition to sleep-like states.

### Estimated model parameters correlate with subjective ratings of sleepiness

The sleep (nap) dataset used in this study contains subjective reports of sleepiness by N=19 participants on two different measures: the Stanford Sleepiness Scale (SSS) and Karolinska Sleepiness Scale (KSS).

In an exploratory manner, we investigated whether estimated parameters from the model (slope and tipping point of sleep drive *α* and *t*_0_, noise *σ*) correlate with any of these characteristics. We found that the pre-nap sleepiness level on the Stanford Sleepiness Scale (SSS) shows a significant positive rank relationship with *α* (Spearman r = 0.65 for 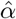 (Fig 8-a) and 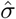 (Fig 8-b); both p-value *<* 0.05*/*3 after Bonferroni correction; correlations with pre-nap KSS scores were consistent, albeit not significant: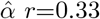, p=.16; 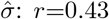, *p*=.07). In other words, participants whose fitted model indicated a steeper sleep drive tended to report higher subjective sleepiness before the nap. One possible, but very exploratory, explanation could be that higher sleep propensity may lead to a more pronounced or rapid transition, although a larger number of participants would be obviously needed to confirm the interpretation. The positive correlation between estimated noise-level and subjective sleepiness before the nap suggests that participants who felt sleepier also exhibited higher intrinsic noise levels, which is seen in Fig. 4 to facilitate earlier and/or more frequent transitions. This potential role of noise, if confirmed, would open an alternative mechanistic view of the wake-to-sleep transition beyond the conventional rising-sleep-drive model, linking internal neural noise to behavior aspects. There was no statistical correlation between model parameters and subsequent KSS/SSS ratings after the nap.

**Fig 8.**
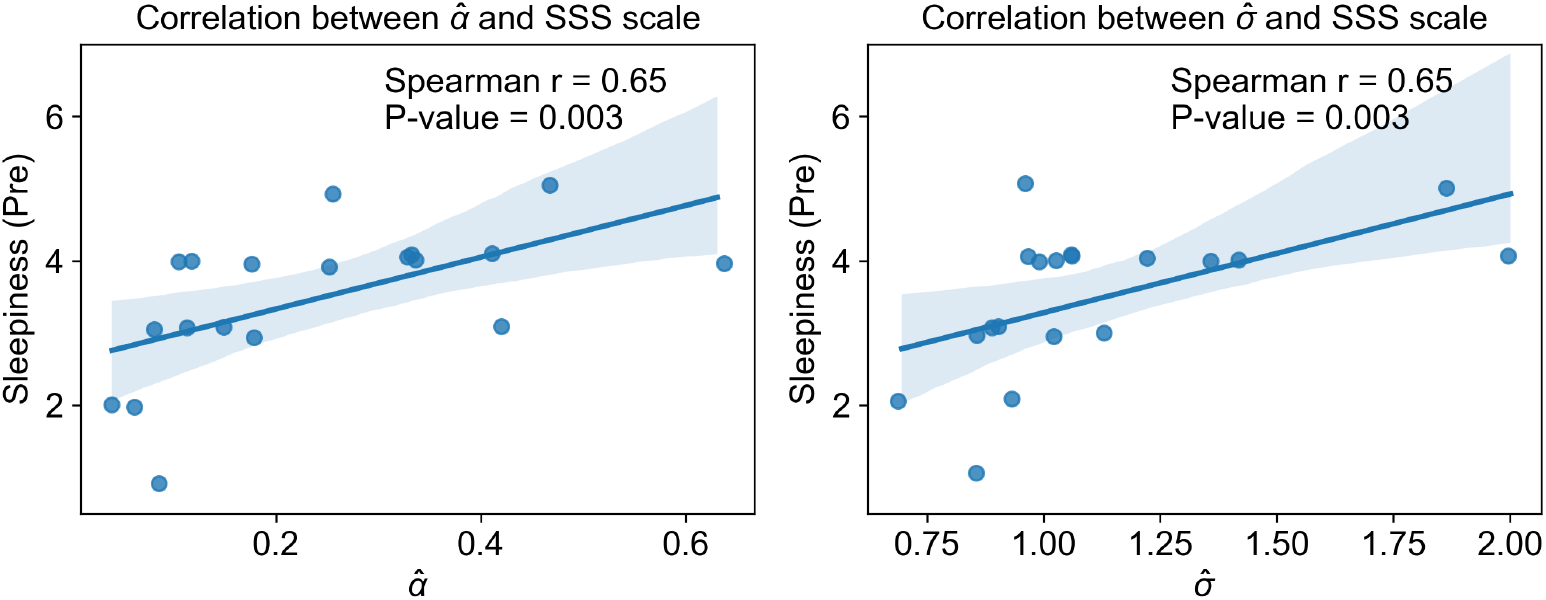
Correlation of fitted model parameters with subjective sleepiness report. **(a-b)** Scatter plots and linear fit model of parameters (left: 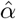, right: 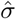) and subjective assessment of pre-nap sleepiness of Stanford Sleepiness Scale (SSS).

**Fig 9.**
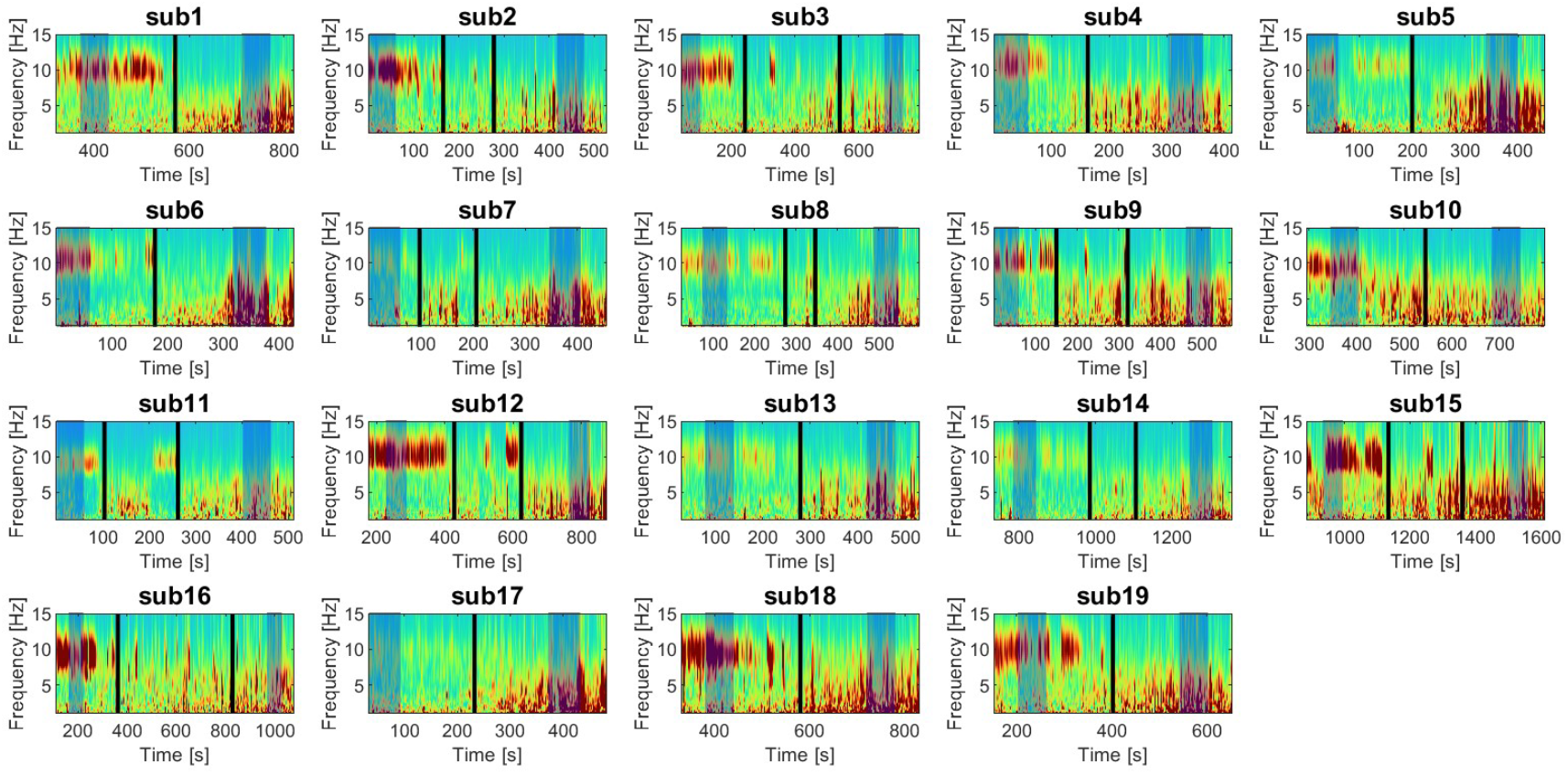
Spectrogram-based validation of SOP windows. EEG wavelet spectrogram for the 19 participants. Each panel corresponds to one subject (**sub1**–**sub19**); the *y* -axis spans 0−15 Hz and the *x* -axis shows time. The two *shaded* regions mark the first one minute (“wake”), and the last one minute (“sleep”), which are used as modes for the embedding. The central, unshaded block is the analysis window. Black vertical bars mark the (possibly identical) time points *t*_*start*_, *t*_*end*_ used to define the analysis window: [*t*_*start*_ - 200s., *t*_*end*_ + 200s.]. Across all individuals, the heuristic ratio amp_*δ*_ */*amp_*α*_ aligns well with the qualitative drop in *α*-power and rise in *θ & δ*-power, confirming the robustness of the SOP definition

## Discussion

### Context and contribution

The sleep-onset period (SOP) is increasingly recognized as a dynamic transition rather than a simple binary switch [11, 12]. Yet traditional analyses, including in the clinic, often reduce it to a single point, typically the first appearance of stage N1 [23]. Even more refined staging methods, such as the Hori classification [24], do not fully encompass the fluctuating continuum during SOP. In response, our study introduces a new modeling framework with three main contributions: (1) a data-driven embedding strategy for high-dimensional EEG time-frequency signals, paired with a parsimonious bistable stochastic model governed by a single, slowly varying parameter that drives the wake-to-sleep transition; (2) a parameter-estimation procedure that is validated in simulated trajectory and real data given single trajectory, thereby allowing systematic quantification of intra- and inter-individual differences in SOP dynamics; and (3) exploratory evidence that estimated model parameters may correlate with subjective ratings of sleepiness around the nap, suggesting a potential utility for differentiating subtypes of sleep-onset difficulties.

Regarding the first contribution, we introduce a parsimonious embedding strategy based on linear interpolating the first SVD mode of wake and sleep states. This embedding not only reduces dimensionality but also preserves key features of the SOP trajectory. Then we model the SOP index (*µ*) as a minimal, parameterized stochastic model in which a slowly changing potential landscape and a tunable noise term together produce a wide range of SOP phenomena, including smooth drift, early or delay switch, as well as noise-driven flickering. Unlike prior models [15], which assume near-equilibrium conditions with negligible noise-level, our approach explicitly incorporates both time-varying parameters (*β*(*t*)) and stochastic fluctuations, allowing us to characterize the interactions between landscape and stochastic forcings during SOP.

Fitting parameters of stochastic model given single trajectory is challenging. Here, we employ a Markov chain Monte Carlo (MCMC) approach to fit its parameters, the slowly varying “sleep drive” and the noise term. We firstly validate the fitting in simulated settings, then apply it on real EEG recordings. The simulated-data experiments reveal that, while certain parameters (e.g., *α, t*_0_) can exhibit moderate variability, their rank ordering across individuals is largely preserved, thus allowing reliable comparisons of inter-individual differences.

With an exploratory correlation analysis, we formulate the hypothesis that model-derived parameters can be potentially served as biomarkers for tracking intra- and inter-individual variability of sleep-onset disorders. Testing this hypothesis will require a large cohort of patients, comparing individuals with conditions like insomnia, delayed sleep phase, or narcolepsy to healthy sleepers. For example, our model may predict that patients with insomnia may have abnormally low or high noise levels, or a slower drift of the sleep drive (*β*), whereas narcolepsy or sleep-deprived individuals might show an abnormally steep drive toward sleep. Such applications could yield quantitative indices for diagnosing and tracking these conditions, complementing existing clinical scales.

### Comparison with previous work

Our findings extend and integrate several threads of prior research on sleep onset dynamics and modeling. Early mechanistic frameworks of sleep-wake regulation (e.g. the two-process model and “flip-flop” switching circuits) established the concept of a bistable control of sleep and wake states, but these models usually involve many variables and parameters with the lack of linking to macroscopic measurements, making them difficult to fit directly to EEG data. On the other end of the spectrum, data-driven approaches have been developed to track sleep onset. For example, Prerau et al. (2014) [25] used a statistical dynamic model to compute a continuous probability of wakefulness by combining EEG and behavioral measures, improving the temporal precision of SOP tracking over traditional sleep-stage scoring, but failed to provide mechanical insight. However, contrary from these approaches, our model explicitly captures the SOP dynamics through a physical description based on stochastic dynamical systems. By gradually adjusting the position of the central barrier between wake and sleep attractors and systematically varying the noise level, our model effectively captures the continuous and stochastic nature of sleep-onset phenomena observed empirically, including intermittent reversals or “flickering” between wake-like and sleep-like states.

### Limitations and future directions

This work has a number of limitations, which we describe here and for which we suggest potential strategies to address them in future work.

#### 1) Parameter estimation in stochastic system

Estimating parameters in stochastic models is challenging. Firstly, a large stochastic perturbation can obscure the underlying system dynamics, making it difficult to accurately estimate the deterministic term. Second, fitting the parameters of a stochastic model to a single trajectory (one SOP per subject) is inherently difficult [26] because the variability observed in one time series may be attributable to multiple distinct parameter sets. Finally, this challenge is amplified in non-stationary systems with time-varying parameters [27]. In future work, we will leverage the neural posterior estimators to match the time-varying, noisy, and single-realization SDE case in our sleep-onset model, which is grounded in recent methodology development of simulation-based inference [28, 29].

#### 2) Embedding strategy and dimensionality

Our current 1-D embedding leverages the idea of mutual inhibition - interpolating between wake and sleep modes (*µU*_*w*_ +(1−*µ*)*U*_*s*_), to capture the rapid switches in EEG spectral content. In practice, this one-dimensional representation, *µ*, could be approximated as an affine transform of *V*_2_ (if one applies SVD across the entire analysis window), given the experimental observation that 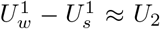 (see Supplementary Materials, where we provide a proof for the affine link between *µ* and *V*_2_). While this consistency validates our simplified linear blending for describing key aspects of the SOP trajectory, it also highlights its inherent limitation: the energy simply shifts from one mode to another mode. Consequently, we miss more nuanced amplitude fluctuations, such as the “flat spectrum”, sometimes observed during transitions and thus fail to accurately reflect other physiological process manifesting as global decreases or gradual shifts in EEG power. In many subjects, this 1-D scheme suffices to capture fundamental wake-sleep transitions; however, going forward, incorporating additional dimensions (e.g., energy fluctuations) approaches would allow the model to accommodate and potentially help uncover additional physiologically relevant process, such as the change of homeostasis.

#### 3) Assumption of one global sleep-drive

Our current model makes a modeling assumption that one global sleep-drive dictates the wake-to-sleep transition, and does not attempt to dissociate the influence of homeostasis and circadian rhythm. Our current formulation of the sleep drive, *β*(*t*), is a single, monotonic, exogenous parameter. While this simplification captures the idea that the overall drive increases during the sleep-onset period, it does not differentiate between distinct physiological components that are known to co-modulate sleep transitions. In neural systems, control signals are often shaped by both a deterministic, feed-forward drive and a state-dependent feedback mechanism. Although the classical circadian rhythm is typically a long-term process, the embedding developed here suggests that during sleep onset, a fast, deterministic component may emerge that prepares the neural system for transition. Without providing a physiologically grounded explanation of what *β*(*t*) represents, our current formulation may oversimplify this complex interplay. In future work, one could aim to disentangle these components, potentially by leveraging dynamic system identification methods (e.g., dynamic SINDy [30]) that can infer separate contributions from feed-forward and state-dependent drives. This refinement will potentially enhance their biological interpretability.

## Materials and methods

### Dataset

N=37 healthy college students from Southwest University in Chongqing (male: 17; female: 20) were enrolled in the study. Inclusion criteria included: no addiction to tobacco or alcohol, no substance abuse, no coffee or other functional drinks in the week before the experiment; no history of neurological or psychiatric diseases; participants needed to have a nap habit (*>* 4 /week and each nap lasting for *>* 30 min); a regular work and rest schedule maintained for 1 week before the experiment (the time to fall asleep no later than 00:00 hours [midnight], the time to wake up between 06:30–08:00 hours, and a total sleep duration of ( 6.5–8 hr); no day–night reversal behaviour; and no crossing time zone behaviour. The subjects completed the Stanford Sleepiness Scale (SSS) and Karolinska Sleepiness Scale (KSS) before nap. Then the subjects had their EEG electronics connected and then took a nap. About 90 min later, the experimenter awoke the subjects and instructed them to complete the scale package (SSS and KSS) one more time, and then again 30 minutes after waking up. Due to a partial lack of sleep scales report, data from N=19 subjects were finally included. The study was approved by the Ethics Committee of the Southwest University.

### EEG recordings

The experiment used a 63 Ag/AgCI electrode cap (Brain Products GmbH, Gilching, Germany) based on the extended 10-20 international electrode position system. Two additional electrodes were used as reference and ground, and the online reference was Fcz The electro-oculogram (EOG) was recorded using two electrodes, one electrode below the left eye and the other outside the tail of the right eye. The EEG signal was recorded with a sampling rate of 500 Hz. Before the experiment, it was ensured that impedance were *<* 5*k*Ω for all electrodes.

The central-occipital electrode (Oz) was selected as it reliably captures the prominent alpha-band (8-12 Hz) activity, then a 0.5 - 30 Hz band-pass filter (4-order Butterworth zero-phase filter) was applied, serving as anti-alias filter and removing high-frequency artifact. After that, the signal was down-sampled down-to 100 Hz. To transform the signal to time-frequency space, we used the analytic (complex) Morlet wavelet (cmor1-1.5: band-width parameter = 1; center-frequency = 1.5 Hz), with defined frequency range of interest to span 0.5 Hz up to 20 Hz, and subdivided that interval into 200 equally-spaced frequency bins. This wavelet-based time-frequnecy representation prepared for the embedding extraction procedure.

### SOP window definition

From each individual participant’s EEG wavelet spectrum, we identified the time window in which the participant transitions from wakefulness to sleep by examining the ratio of delta-band (0.5–4 Hz) *amp*_*δ*_ to alpha-band (8–12 Hz) amplitudes *amp*_*α*_. The start of transition (*t*_*start*_) was defined as the earliest continuous block of time where 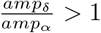 for more than one minute. The end of transition (*t*_*end*_) was identified as the earliest continuous block of time where 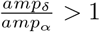 for more than two minutes. We then defined the SOP window as [*t*_*start*_ − 200*s*., *t*_*end*_ + 200*s*.]. Within this window, we labelled the first 1 minute as “wake” and the last 1 minute as “sleep”. Although this definition is entirely heuristic, we used it consistently across all individuals. All individual embeddings can be found in the (Fig 2-right).

### Embedding extraction

Using the SOP window, we then construct a generalized, low-dimensional representation *µ*(*t*) of the SOP spectrogram by computing the singular value decomposition (SVD) of the time-frequency (TF) representation separately in the “wake” and “sleep” segments to obtain the first principal modes, 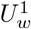 and 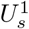. We then generate the embedding *µ*(*t*) by projecting the spectrogram on 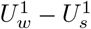. To do so, we first normalized each spectrogram bin in the SOP window to the unit norm, then projected onto 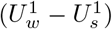 to obtain a scalar *µ*(*t*). To scale the *µ*(*t*) to the same range of model output, a linear transform (2 *× µ*(*t*) − 1) was applied. Finally, we applied a mild gaussian-window smoothing and down-sampling by a factor of ten to discard high-frequency noise that the model not intend to explain while improving the computational efficiency.

Conversely, to project state-space dynamics back into observation (EEG) space, we reconstruct a low-rank spectrogram using the linear interpolation 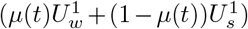 (Fig 2a-bottom).

### Model Structure

To model the low-dimensional dynamics *µ*(*t*), we propose a minimally-parameterized first-order model of the form given in Eq. 2, which we reproduce here for convenience:

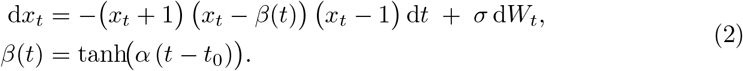

The system has by construction two stable fixed points at *x* = *±*1 and a single unstable fixed point whose location is determined by the time-varying parameter *β*(*t*). *β*(*t*) is modeled as a slowly-drifting hyperbolic tangent function, with *α* controlling the speed of its changing rate, and *t*_0_ dictating its tipping point.

### Parameter Inference

The macroscopic embedding we extracted *µ*_0:*N*_ is assumed to be the latent trajectory generated from stochastic dynamical system (SDE) corrupted by additive gaussian observation noise:

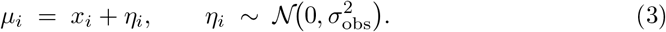

The latent state *x*_0:*N*_ itself evolves according to the cubic SDE (Eq. 2), with drift parameters *θ*_*drift*_ = (*α, t*_0_) and diffusion parameter *θ*_*diff*_ = *σ*. The EEG recordings and the simulator don’t necessarily share the same physical clock: the effective sampling may be slower or faster than the nominal integration steps used in the SDE step. We therefore introduce a time-scale factor, *t*^⋆^ = *ε t*, which stretches (*ε >* 1) or compresses ((*ε <* 1) the model time axis so that the latent state can align with an unknown true time-scale of experimental data. Hence, all unknowns are therefore collected in Eq. 4:

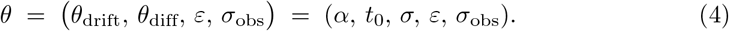

According to Bayes’ rule, the joint posterior density function over parameters and latent path is proportional to

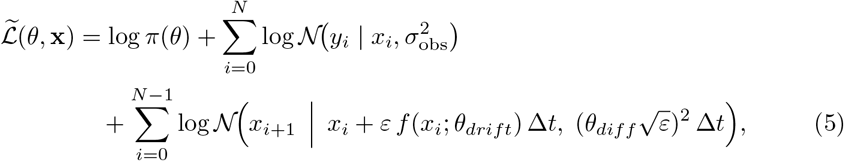

where *π*(*θ*) is the joint prior, and *f* (*x*_*i*_; *θ*_*drift*_) = −(*x*_*i*_ − 1)(*x* − *β*_*i*_)(*x*_*i*_ + 1), *β*_*i*_ = *tanh*(*αε*(*t*_*i*_ − *t*_0_))

The three parameters we are interested in are {*α, t*_0_, *σ* }. For *α*, its prior is set to a half-normal distribution with unit standard deviation (*α*_prior_ ∼𝒩^+^(0, 1)). This forces the fitting procedure to emphasize small alpha regimes, for the two following reasons: first, the timescale of hyperbolic tangent function is inversely proportional to *α*, which means its changing rate is mostly sensitive to the change of *α* when it’s small, and unidentifiable for large *α* (Fig 6-d). Second, it’s a reasonable assumption that the slow drive (a combination of homeostatic pressure and circadian rhythm) evolves in a relatively slow scale. The prior of *t*_0_ is set as normal distribution, 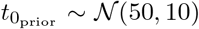 (simulated case) or 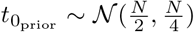 (experimental data). Only three *t*_0_ values were tested in simulated case, because, first, defining highly standardized initial time across subjects is difficult in practical real-data settings and, second, increasing the number of distinct *t*_0_ values further increases a lot of computational cost by requiring more loops. The prior distribution of *σ* was *σ*_prior_ ∼𝒩^+^(0, 2) in both simulated and real data to force MCMC only sample the positive values with a relatively broad range. The prior of the time-scale factor, *ε*, is set as a uniform distribution, *ε*_prior_ ∼ 𝒰 (0.1, 5), which allows the fitted dynamics to be up to ten times slower (*ε* = 0.1) or five times faster (*ε* = 5) than the EEG step size. The lower bound prevents numerical stiffness, while the upper bound still covers all physiologically plausible transition speeds encountered across individuals.For observational noise, we assumed a Gaussian observation noise: *σ*_*obs*_ ∼ 𝒩 (*µ*, std), where *µ* is given by the latent state simulated from SDE. In the simulated case, std is set as 0.00001 to force the latent state trajectory simulated in MCMC to match the observed simulated trajectory almost exactly. In the real data case, std ∼𝒩 (0.1, 0.4) to allow the latent states to differ from the observed data and adapt to potentially different noise-levels to different individuals. The detailed parameter settings can be found in Tables 1–2 (simulated data) and Table 3 (experimental data).

**Table 1.** Fixed–landscape scenario: simulation settings and priors.

| Specification | Value(s) | Prior |
| --- | --- | --- |
| Number of trajectories | 30 | — |
| Initial state $X_0$ | 0 | — |
| Unstable fixed-point location $\beta$ | $[-0.8:0.2:0.8]$ | $\mathcal{N}(0, 1)$ |
| Noise $\sigma$ | $\{0.2, 0.5, 1.0\}$ | $\mathcal{N}^+(0, 2)$ |

**Table 2.**
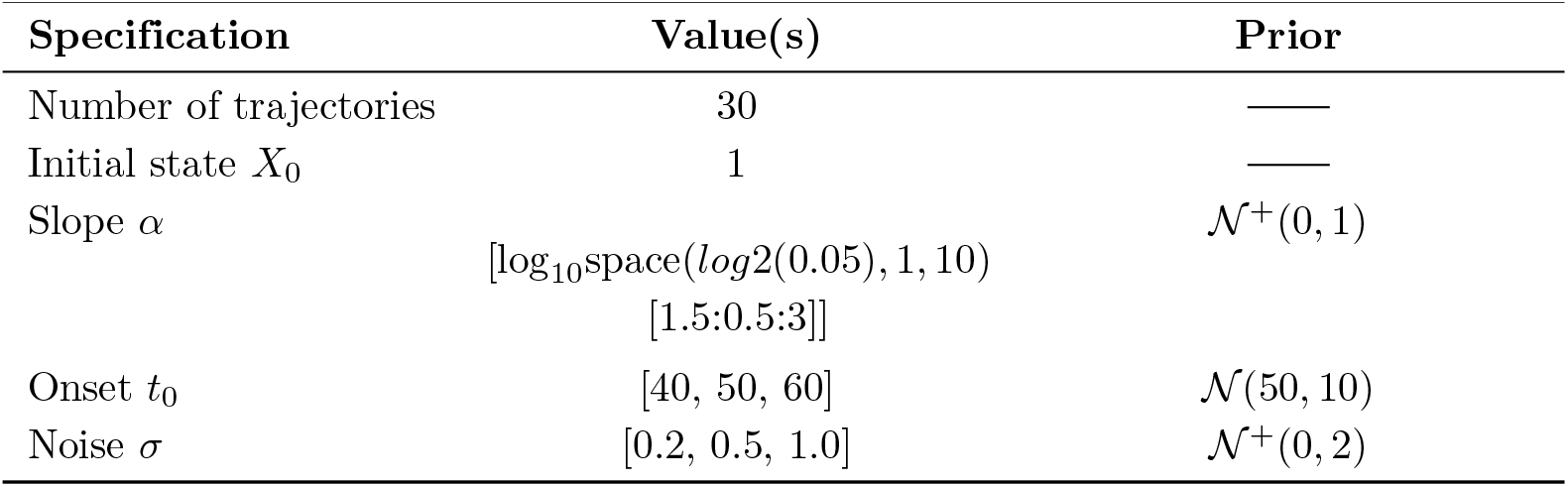
Time-varying landscape scenario: simulation settings and priors.

**Table 3.** Priors adopted for real EEG data.

| Parameters | Prior |
| --- | --- |
| $\alpha$ | $\mathcal{N}^+(0, 1)$ |
| $t_0$ | $\mathcal{N}(\frac{N}{2}, \frac{N}{4})$ |
| $\sigma$ | $\mathcal{N}^+(0, 2)$ |
| $\varepsilon$ | $\mathcal{U}(0.1, 5)$ |
| $\sigma_{\text{obs}}$ | $\mathcal{N}^+(0.1, 0.4)$ |

### Model evaluation

In simulated settings, we use Spearman’s rank correlation coefficient to evaluate the fitting performance for *α*, as the goal is not accurately capturing the true value, but consistently differentiating its order.

For real EEG data, since we don’t have access of the true value, and model is stochastic, wee applied posterior predictive check to compare between what the fitted model predicts and the actual observed data via running forward simulation of 4000 trajectories from the joint posterior distribution and selected the one with minimal Root Mean Square Error (RMSE) value as well as the one maximizing joint posterior density, then plot its time course and corresponding reconstructed time-frequency plot to compare with the SVD embedding, taken as ground truth.

## Acknowledgments

Work funded by the MSCA Doctoral Network LULLABYTE (to JJA, ZH), Region Bourgogne Franche-Comté ANER action ASPECT (to JJA, MA), National Natural Science Foundation of China 32471095 (to XL). Work conducted in the framework of the EIPHI Graduate school (ANR-17-EURE-0002 contract). The work of JNK was supported in part by the US National Science Foundation (NSF) AI Institute for Dynamical Systems (dynamicsai.org), grant 2112085. JNK further acknowledges support from the Air Force Office of Scientific Research (FA9550-24-1-0141). We thank Björn Rasch, Thomas Andrillon, Delphine Oudiette, Megan Morrison, Lou Zonca and Richard Gao for useful discussions on previous versions of this work.

## Supplemental Material

### A Validation of the SOP window across 19 subjects

For every participant, we defined a *Sleep Onset Period* (SOP) by detecting the transition from wakefulness to sleep in the continuous EEG spectrogram, following the heuristics described in the *Methods* section.

### B Affine link between *µ*(*t*) and *V*_2_

#### Notation and dimensions

- *M* ∈ ℝ^*F×T*^ — raw spectrogram (*F* frequency bins, *T* time points);
- *D* ∈ ℝ^*T ×T*^ — diagonal, 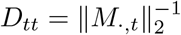 (each column of *M* is rescaled to unit 𝓁_2_ norm);
- 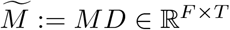 — *normalised* spectrogram;
- *U* ∈ ℝ^*F×F*^ , *V* ∈ ℝ^*T ×T*^ , Σ = diag(*σ*_1_, *σ*_2_, … ) — singular-value decomposition (SVD) of the *raw* matrix *M* :

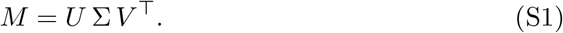
- 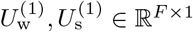 — first left singular vectors estimated on wake-only and sleep-only segments;
- 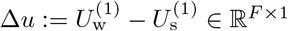.

#### Projection that defines *µ*(*t*)

For every column of the *normalised* spectrogram we project onto the direction Δ*u*:

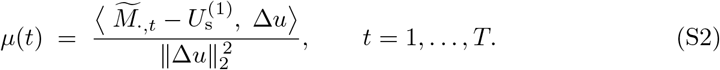

Gathering the scalars into a row vector *µ*^⊤^ = [*µ*(1), … , *µ*(*T* )] ∈ ℝ^1*×T*^ gives

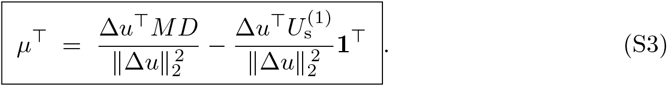

#### Empirical rank-2 structure

Fig. 10 shows that the second frequency mode of *M* coincides with Δ*u* for most subjects:

**Fig 10.**
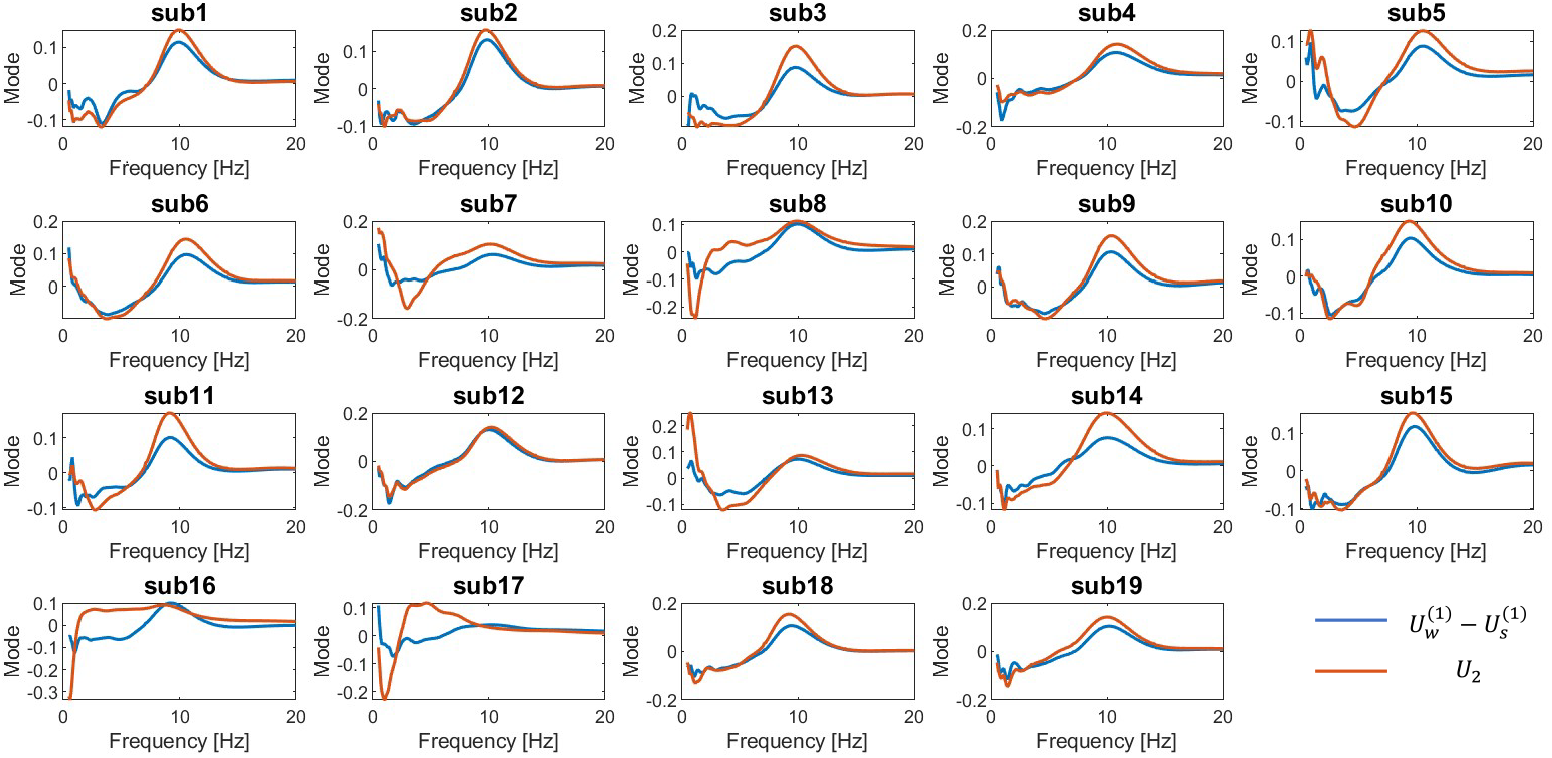
Comparison between SVD modes. For every participant, the difference between the first left singular vectors extracted from wake-only 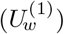 and sleep-only 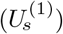 spectrogram blocks (blue) closely matches the second left singular vector of the full raw spectrogram, *U*_2_ (orange).

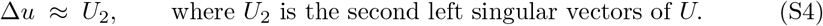

#### Action of 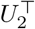 on the *MD* product

Left-multiplying (S1) by 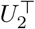 and right-multiplying by *D* yields

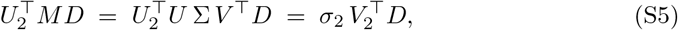

#### Affine relation between *µ* and *V*_2_

Substituting (S4) into (S3) and then invoking (S5) gives the *explicit* affine map

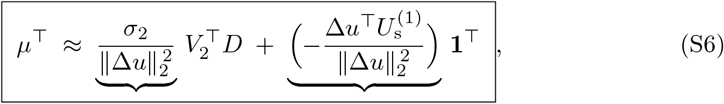

Equation (S6) confirms that the one-dimensional embedding *µ*(*t*) and the right singular vector *V*_2_ (after the column-wise normalisation encoded by *D*) contain the *same* information up to a fixed scaling *a* and offset *b*.

### C Posterior predictive analysis: minimal-RMSE trajectory and KL/RMSE summaries

**Fig 11.**
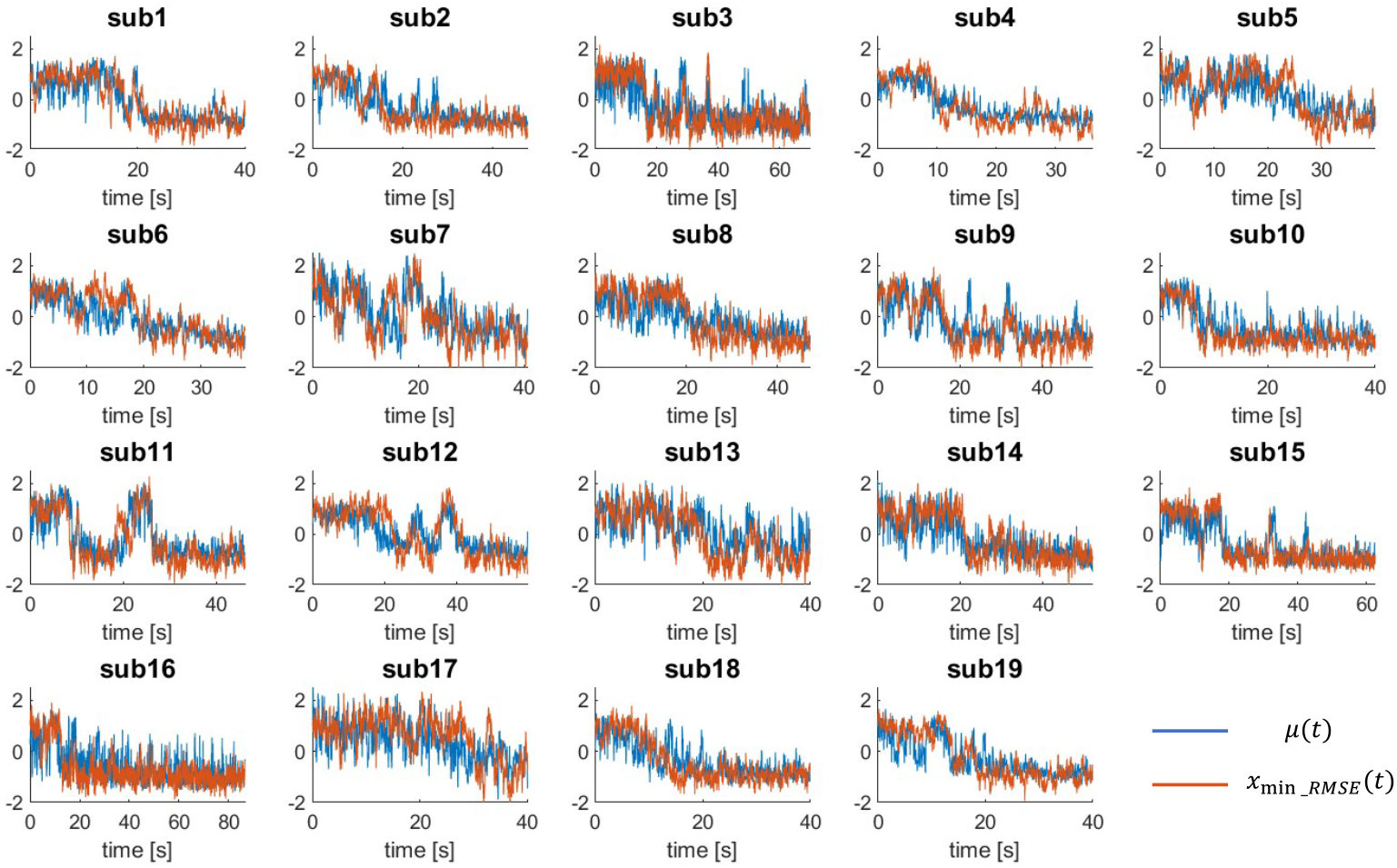
Example posterior draw with minimal RMSE. For each subject we ran a MCMC inference over the model parameters and generated 4000 posterior predictive trajectories *x*_sim_(*t*) from the joint-posterior distribution. The orange line in every panel shows the simulated trajectory with the smallest root-mean-square error (RMSE) to the embedding *µ*(*t*) (blue)

**Fig 12.**
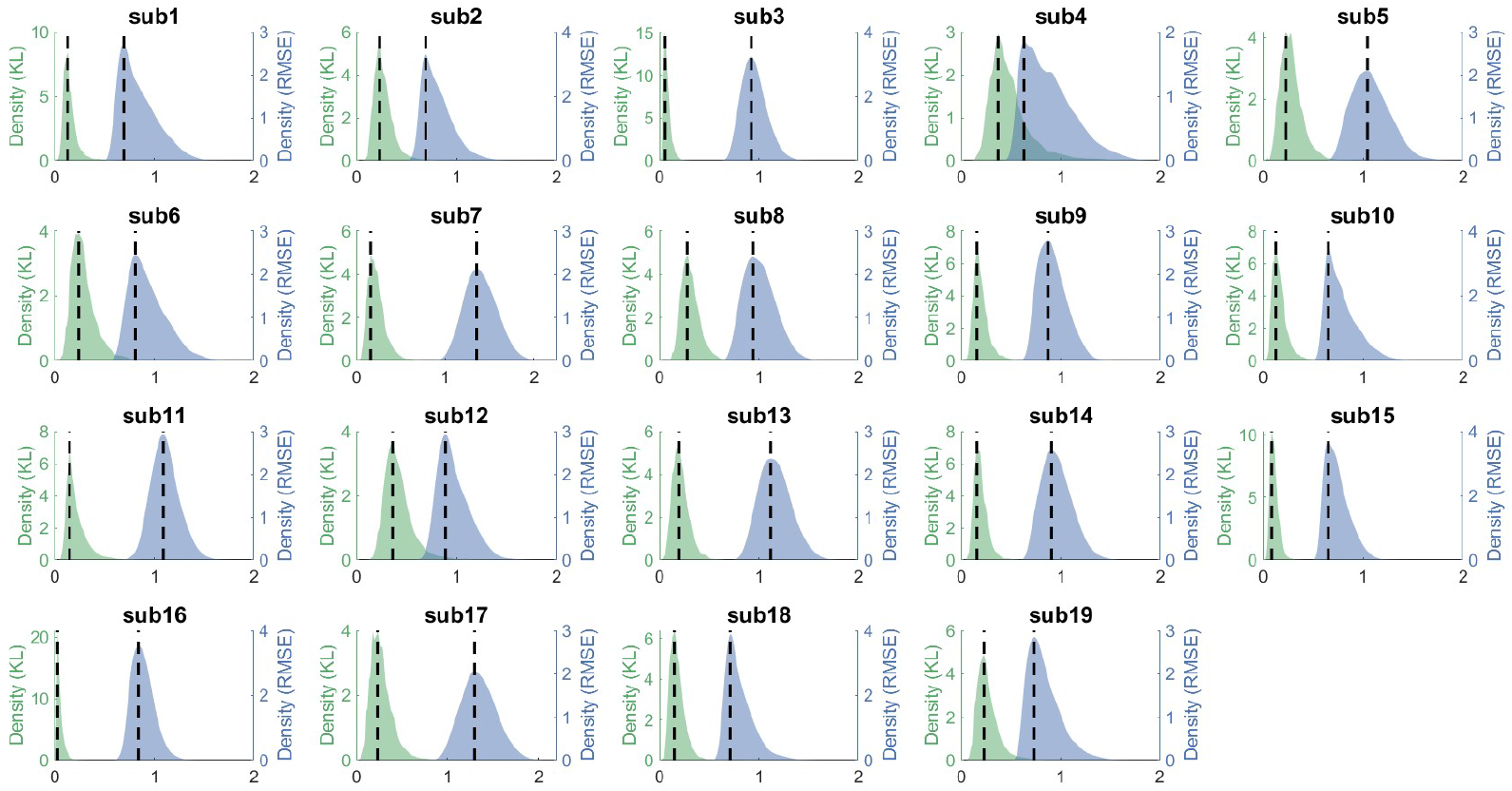
Distribution of KL-divergence and RMSE over posterior samples. Posterior predictive checks were performed by forward simulation of 4000 trajectories from joint posterior distribution per participant and computing two complementary metrics: Kullback–Leibler (KL) divergence between the *µ*(*t*) and simulated trajectories (green, *left* axes) and point-wise RMSE (blue,*right* axes). Dashed vertical lines indicate the median of each distribution. For all subjects the bulk of the KL values lie below 0.16 and RMSE clusters around 0.8, confirming that a large portion of parameter draws generates trajectories statistically close to the data. Together with the illustrative trajectories in Fig. 11, these metrics demonstrate the overall adequacy of the inferred model.

## Footnotes

1 by phenomenological, we refer here to models that attempt to represent observable properties of their targets without necessarily being derivable from an existing underlying theory [5]. In neuroscience and neurology, the word ‘phenomenological’ is also used to relate to a participant’s subjective experience, but this is not the meaning we use here.

